# Molecular mechanisms behind global distribution of earthworm revealed by the genome

**DOI:** 10.1101/853267

**Authors:** Xing Wang, Yi Zhang, Yufeng Zhang, Mingming Kang, Yuanbo Li, Samuel W James, Yang Yang, Yanmeng Bi, Hao Jiang, Xin Zhou, Yi Zhao, Zhenjun Sun

**Author notes:** These authors contributed equally: Xing Wang, Yi Zhang, Yufeng Zhang, Mingming Kang. Correspondence to: College of Resources and Environmental Sciences, China Agricultural University, Beijing 100193, China E-mail address (X. Wang); (Y. Zhao); (Z.J. Sun).

## Abstract

Earthworms (Annelida: Crassiclitellata), are widely distributed around the world due to their great adaptability. However, lack of a high-quality genome sequence prevents gaining the many insights into physiology, phylogeny, and genome evolution that could come from a good earthworm genome. Herein, we report a complete genome assembly of the earthworm *Amynthas corticis* of about 1.2 Gb, based on a strategy combining third-generation long-read sequencing and Hi-C mapping. A total of 29,256 protein-coding genes are annotated in this genome. Analysis of resequencing data indicates that this earthworm is a triploid species. Furthermore, gene family evolution analysis shows that comprehensive expansion of gene families in the earthworm genome has produced more defensive functions compared with other species in Annelida. Quantitative proteomic iTRAQ analysis shows 97 immune related proteins and 16S rDNA sequences shows 88 microbes with significantly response to pathogenic *Escherichia coli* O157:H7. Our genome assembly provides abundant and valuable resources for the earthworm research community, serving as a first step toward uncovering the mysteries of this species, may explain its powerful defensive functions adapt to complex environment and invasion from molecular level.

## Introduction

The earthworm belongs to oligochaete annelid distributed all over the world. Recently, a research focus on diversity of earthworm at the global scale showed that this underground species has more local richness and abundance in temperate zone, but more dissimilarity across different locations in tropical zone^1^. More interestingly, it also showed that climate is the key factor affecting distribution of earthworm, in spite of the soil property and habitat cover through modeling based on integrated big data^1^. Soil properties and habitat covers represent diverse environments, which means different threats and dangers to species, especially for underground ones. Thus, it could be proposed that there must be unique and robust molecular mechanisms in individual earthworm helping it face to and live in adverse and complex environments.

Earthworm has been recognized as the species with profound ecological and economic impacts on soil^2,3^ and considered as the “soil engineer”. On the other hand, earthworm is also an invasive species with not ignorable affections on soil ecosystem^4-6^, including local diversity of native species^7^. Behind the invasive characteristic, a close association between polyploidy and parthenogenetic reproduction has been identified in earthworm species of *Lumbricidae* family^8-11^. Polyploidy of earthworm provides more genomic materials for evolving novel phenotypes, while parthenogenetic reproduction ensures the fixation of them, which is benefit for its rapid adaptation to a new environment.

*Amynthas corticis*, an earthworm species of *Megascolecidae* family, which is a wild species with global distribution. Particularly, *A. corticis* is also a cosmopolitan invasive species originating from Asian. There was a study suggested that the invasive ability of *A. corticis* is due to its greater mobility and polyploidy or parthenogenetic reproduction^12^. To reveal the whole picture of molecular mechanisms behind the strongly adaptive ability of *A. corticis*, we sequenced its genome and generated a complete assembly with a length of 1.29 Gb in total, including 42 chromosome-level scaffolds with N50 length of 31 Mb (Table 1 and Supplementary Fig. 1). In addition to the genome, we also detected protein mass spectrum for the body of earthworm and sequenced 16S rDNA for the intestinal tract of earthworm under the treatment of a pathogenic *Escherichia coli* stress *O157:H7*^*13*^, which mimics the severely adverse environmental factor, to profile how earthworm genome functions when facing to stresses.

**Table 1.** Summary of features related to genome assembly.

| Assembly feature | Contig | Scaffold |
| --- | --- | --- |
| Total length (bp) | 1,190,923,171 | 1,285,148,884 |
| Number of sequences | 16,882 | 1,042 |
| Length of the longest sequence (bp) | 1,225,102 | 55,651,430 |
| N50 length (bp) | 117,165 | 31,950,757 |
| N90 length (bp) | 33,690 | 21,414,444 |
| GC content | 40.34% | 40.34% |
| N content | 0 | 0.26% |
| Number of gaps | 0 | 12,771 |
| Length of gaps (bp) | 0 | 3,283,278 |

## Results

### *A. corticis* is triploid

Based on the data resequenced from two individual earthworms of *A. corticis*, we called genome wide SNPs and constructed the coverage distribution of heterozygous k-mers. Following that, we evaluated the ploidy of the genome of *A. corticis*. First of all, the frequency distribution of SNPs at biallelic loci showed that there were two peaks near the frequency of 1/3 and 2/3 appeared (Fig. 1d), suggesting most of biallelic loci have three copies in the genome of *A. corticis*. In addition, the result of statistical analysis based on the frequency distribution of genome wide SNPs showed the feature of triploidy possesses the lowest delta log-likelihood (Fig. 1c), suggesting the genome of *A. corticis* is triploid with the largest probability. Furthermore, the coverage distribution related to 88% of heterozygous k-mers concentrated at 3n for the total coverage of k-mer pairs and 1/3 for the normalized coverage of minor k-mers (Fig. 1e), implying the triploid feature of the genome of *A. corticis*. Lastly, karyotype analysis of four earthworms showed that chromosome number of each individual was larger than 120 (Fig. 1a-b). In combination with that we have assembled 42 chromosome-level scaffolds, the large number of chromosomes also implied the triploid feature of *A. cortices* genome. Thus, multiple independent proofs supported the genome of *A. corticis* is triploid.

**Figure 1.**
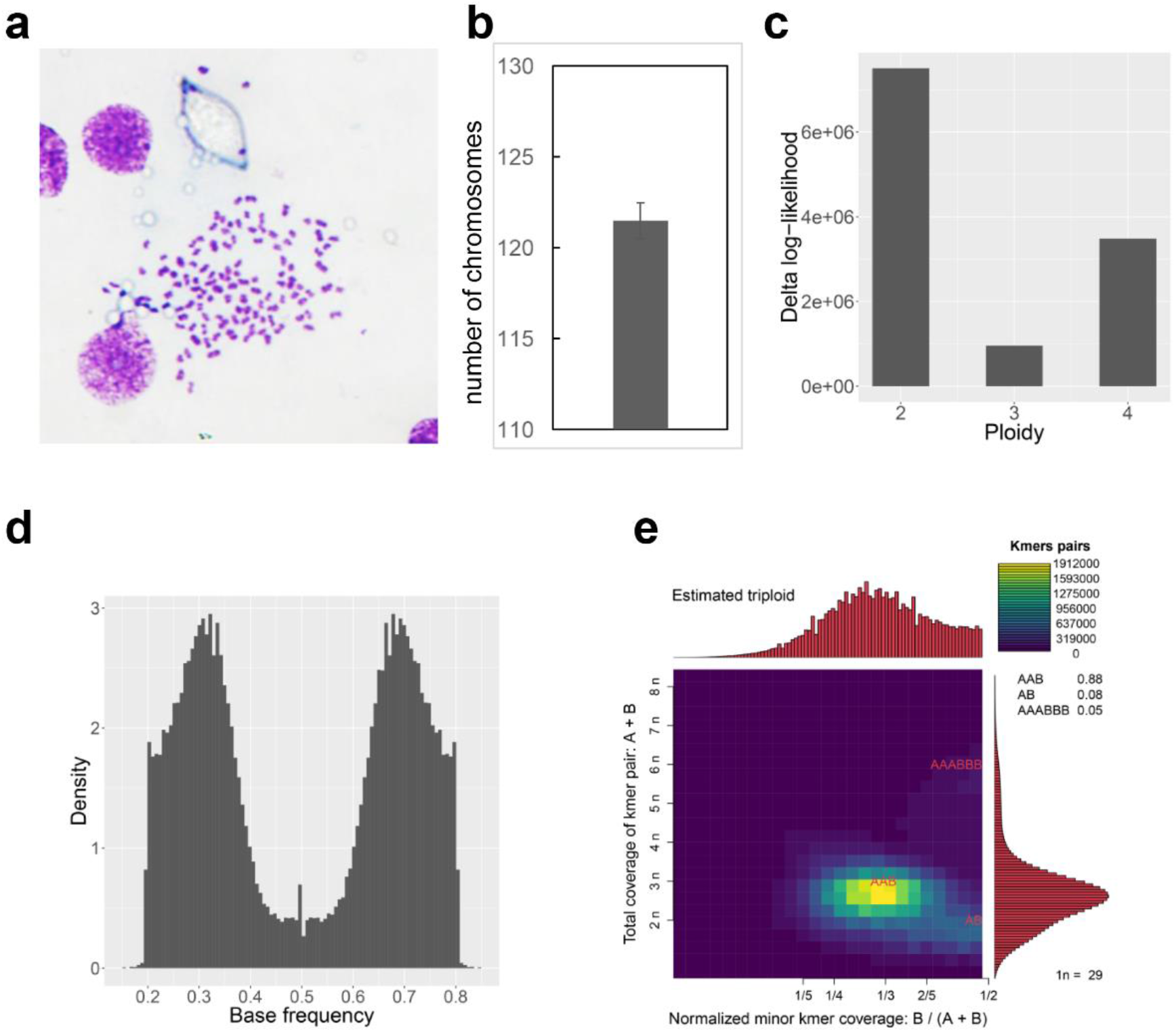
The triploidy feature of *A. corticis*. **a**, Imaging of chromosomes in an individual *A. corticis* resulted from karyotype analysis. **b**, Chromosome numbers of four *A. corticis* individuals determined via karyotype analysis. **c**, Delta log-likelihood values under hypotheses with different ploidy features. **d**, Distribution of frequencies related to biallelic loci. **e**, Coverage pattern of heterozygous k-mer pairs, in which X axis indicates normalized minor k-mer coverage, while Y axis indicates the total coverage of heterozygous k-mer pairs.

### Abundant gene contents related to defensive

Compared with other two annelids, *Capitella teleta* and *Helobdella robusta*^14^, the genome size of earthworm *A. corticis* is significantly larger than them with 3.87 and 5.49 folds, respectively. This phenomenon suggests that there should be comprehensive expansion of gene families happened in earthworm genome. To make it clear, we constructed orthologous groups for genes in earthworm and other invertebrate lineages, including annelids, mollusc, flatworms, roundworms and vertebrates based on TreeFam database^15,16^.

To reveal the evolutionary relationship of earthworm and other species, we construct the phylogenetic tree covering all of species mentioned above based on single-copy orthologous groups. As is shown in Figure 2a, the constructed tree revealed evolutionary relationship of all species consistent with the common sense. However, when focusing on orthologous groups with significantly evolutionary rate, the Spearman’s rank correlation coefficient matrix of group size of different species shows earthworm was away from its phylogenetic tree neighbor of annelids and mollusc, but close to flatworms, roundworms and two snakes in vertebrates (Fig. 2b). These contradictory phenomena imply that environmental or other factors had affected the evolutionary path of earthworm away from other annelids, resulting in a unique annelid genome.

**Figure 2.**
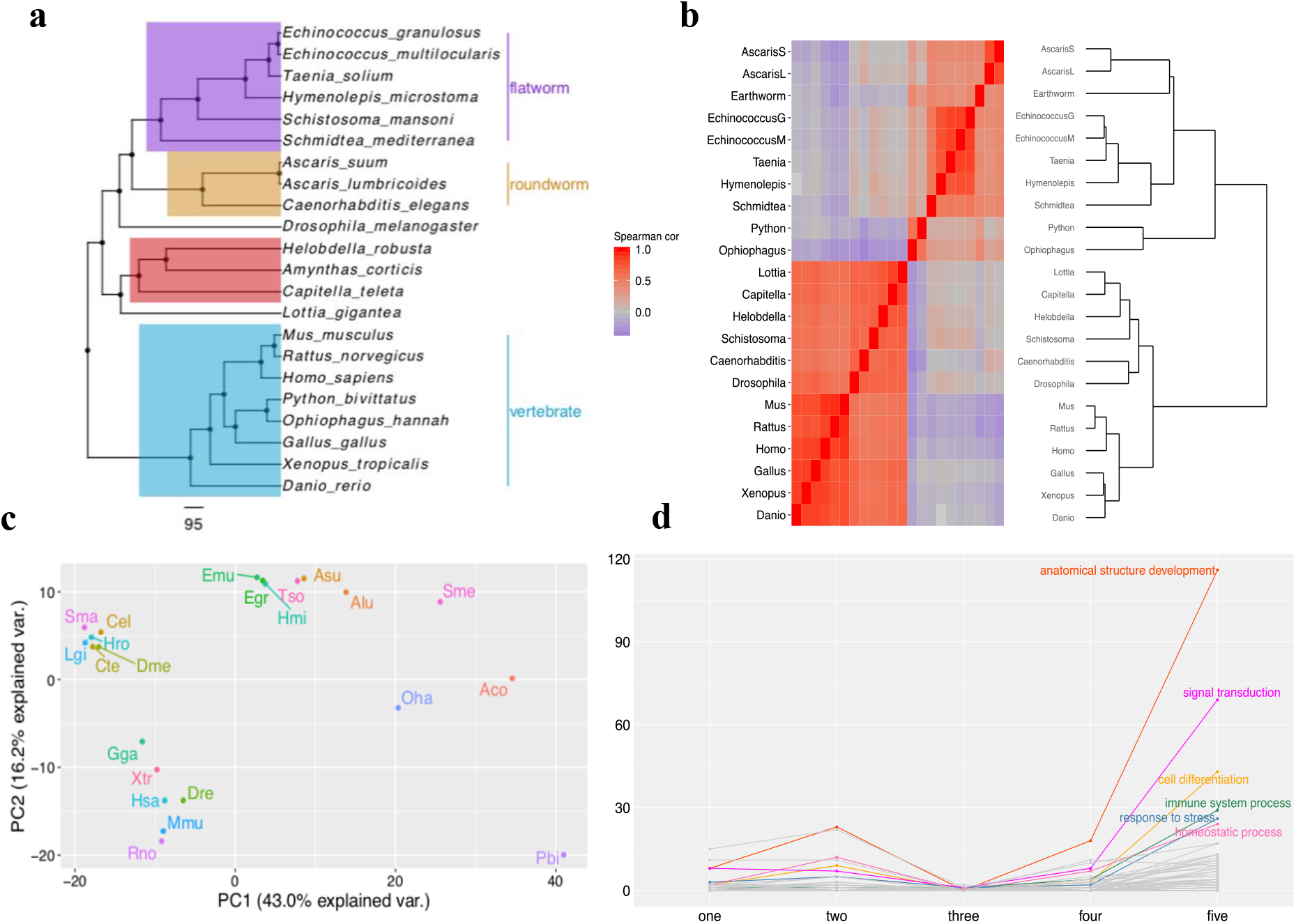
Species tree and evolving pattern showed by orthologous groups with significantly evolutionary rate. **a**, Species tree constructed based on conserved single-copy orthologous groups. The scale indicates 95 million years. **b**, Heatmap and dendrogram of Spearman’s rank correlation coefficient matrix between each pair of species related to gene counts in orthologous groups with significantly evolutionary rate. The distance between each pair of species was defined as the difference value between 1 and Spearman’s rank correlation coefficient. **c**, Clustering of species revealed by principal component analysis (PCA) based on gene counts in orthologous groups with significantly evolutionary rate. **d**, Count of enriched functions at each evolutionary step, in which each line represents a category of GO slims. AscarisL or Alu: *Ascaris lumbricoides*, AscarisS or Asu: *Ascaris suum*, Caenorhabditis or Cel: *Caenorhabditis elegans*, Capitella or Cte: *Capitella teleta*, Danio or Dre: *Danio rerio*, Drosophila or Dme: *Drosophila melanogaster*, Earthworm or Aco: *Amynthas corticis*, EchinococcusG or Egr: *Echinococcus granulosus*, EchinococcusM or Emu: *Echinococcus multilocularis*, Gallus or Gga: *Gallus gallus*, Helobdella or Hro: *Helobdella robusta*, Homo or Hsa: *Homo sapiens*, Hymenolepis or Hmi: *Hymenolepis microstoma*, Lottia or Lgi: *Lottia gigantea*, Mus or Mmu: *Mus musculus*, Ophiophagus or Oha: *Ophiophagus hannah*, Python or Pbi: *Python bivittatus*, Rattus or Rno: *Rattus norvegicus*, Schistosoma or Sma: *Schistosoma mansoni*, Schmidtea or Sme: *Schmidtea mediterranea*, Taenia or Tso: *Taenia solium*, Xenopus or Xtr: *Xenopus tropicalis*.

Detailed analysis of orthologous groups with significantly rapid evolutionary rate showed that more than 70% of them in earthworm are expanded compared with the most recent common ancestor (MRCA) of annelids, which is consistent with its large genome size. Furthermore, the principle component analysis (PCA) using group size of different species also showed that earthworm is away from other annelids along the first principle component (Fig. 2c). What is more interesting, earthworm diverged away from other annelids with a stepwise manner. At each step, species of flatworms, roundworms or vertebrates appeared. For convenience, we defined five steps along the divergence, which are represented by *Echinococcus multilocularis, Ascaris lumbricoides, Ophiophagus Hannah, Schmidtea mediterranea* and itself. Then we extracted orthologous groups with the size in earthworm, which is larger than it in annelids and mollusc while smaller than or equal to it in representative species of each step, as expanded gene families of earthworm at this step. The result of functional enrichment analysis^17,18^ showed that there were the most enriched expanded functions at the last step. Besides fundamental functions such as anatomical structure development, signal transduction, cell differentiation and transport, many functions related to immune system process, response to stress and homeostatic process expanded (Fig. 2d). It could be suggested that the evolutionary process at the last step ensures abundant gene contents related to defensive in the genome of *A. corticis* and thus equips earthworm with large number of molecular arms and effective strategies to live in hostile environments. Furthermore, we reconstructed the evolutionary path of functions related to immune system process, response to stress and homeostatic process through plotting odds ratio at each step for significantly enriched ones (Supplementary Fig. 2-4).

### Defensive to pathogenic *E. coli O157:H7*

We fed earthworm with pathogenic *E. coli O157:H7*^*13*^, which mimics the severely adverse environmental factor, and then detected protein mass spectrum for the body of earthworm and sequenced 16S rDNA for the intestinal tract of earthworm. To profile the dynamic pattern of expression and intestinal microorganism constitution, we selected samples at several time points after treatment of pathogenic *E. coli O157:H7* to detect protein mass spectrum while sequence 16S rDNA. Following normalization of expression level and 16S rDNA abundance, we conducted analysis of variance (ANOVA) for both of them.

In terms of protein mass spectrum, we found 23 genes expressed with significant variation among samples (Fig. 3). First of all, none of these 23 genes expressed in control samples, indicating their specific roles in response to stresses. Furthermore, these genes could be classified as two categories according to whether expressed on the immediate 3^rd^ day after feed of pathogenic *E. coli O157:H7*. Genes expressed on the 7^th^ day or both of the 7^th^ and 28^th^ days but not in the 3^rd^ day were classified as the first category, while genes expressed on the 3^rd^ day were classified as the second category. There are 6 genes belonging to the first category, representing the pattern of later regulation to resist pathogenic bacteria. Two of these genes contained abhydrolase domain or F-BAR domain, while others were novel ones appeared in earthworm without any reliable functional counterparts in otherwise species. The left 17 genes belong to the second category, representing a pattern that immediate activation followed by subtle regulation. Genes classified as the second category include heat shock 70kDa proteins, M1 metallopeptidases, and SRCAP complex, which expressed not only on the 3^rd^ day but on either the 7^th^ or 28^th^ day.

**Figure 3.**
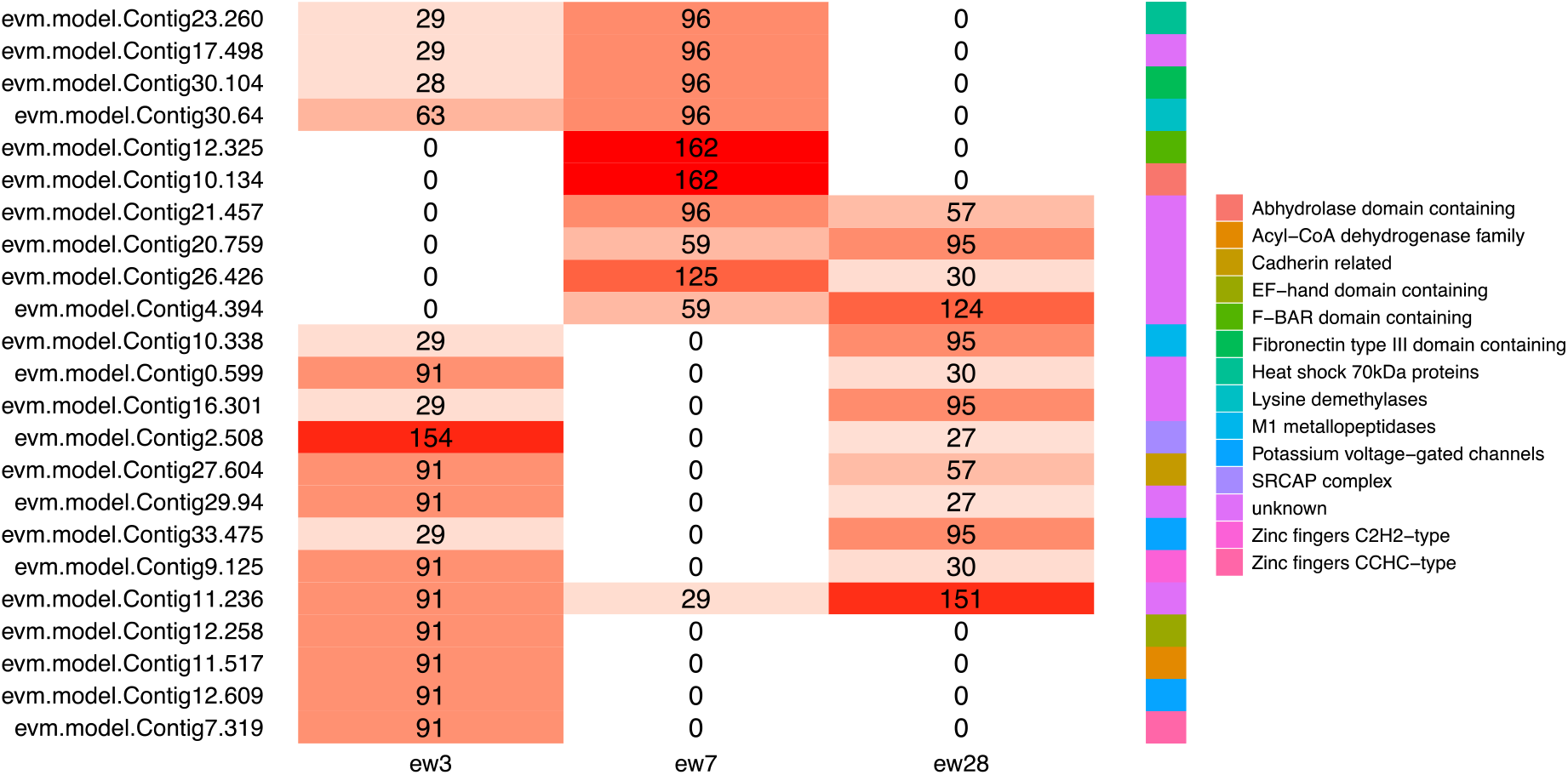
Expression heatmap of earthworm genes with significantly differential expression along days after feeding of pathogenic bacteria. In which, normalized expression count produced by protein mass spectrum and gene annotation retrieved from HGNC database were labeled.

For 16S rDNA, we found the abundance of 9 kinds of microorganisms varied significantly among samples (Fig. 4). The dynamic changing patterns of abundance of these 9 kinds of microorganisms could be classified as 2 categories, including down-up trend and novel appearance on the 28^th^ day. The pattern of down-up trend revealed the reconstruction process of native microorganisms in the intestinal track of earthworm, which could be considered as an indicator representing defensive effects to the pathogenic bacteria. In particular, soil microorganisms belong to class *Acidobacteria-6*^19^ or order *Ellin6067*^20^ reached the highest abundance on the 28^th^ day compared with others, revealing the active uptake of earthworm to resist biotic stresses. Microorganisms with novel appearance on the 28^th^ day, including drilosphere microorganisms belong to family *AKIW874*^21^, also revealed the active uptake of earthworm helping to survive under stresses.

**Figure 4.**
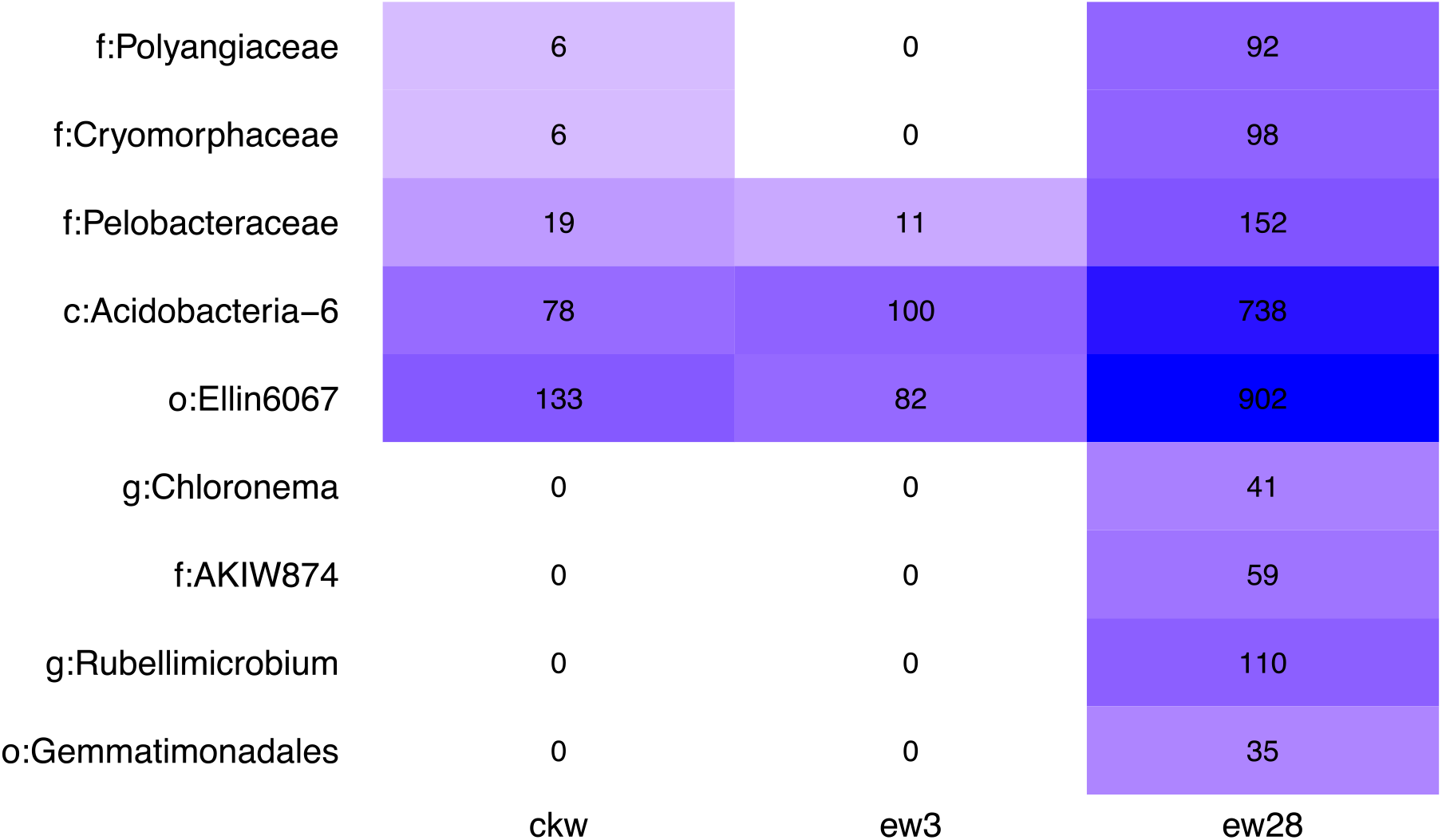
Abundance of microorganisms in intestinal tract of earthworm with significant difference along days after feeding of pathogenic bacteria. In which, normalized abundance and species origin of microorganisms were labeled.

### Profile of defensive genes in earthworm genome

Great adaptation and strong vitality of earthworm have always attracted attentions of biologists and ecologists to make it clear. Thus, several well determined defensive genes were cloned and studied in detail^22-37^. However, due to the absence of a complete genome, it is failed to construct an overview on defensive systems of earthworm. Here, we systematically profiled copies of defensive genes with reliable evidence in earthworm from the assembled complete genome. Predicted genes were considered as orthologs of well determined defensive genes if there were reciprocally best hits between them, which were listed in Supplementary Table 1. Then, we analyzed genomic location and expression pattern for orthologous group members of defensive genes (Fig. 5).

**Figure 5.**
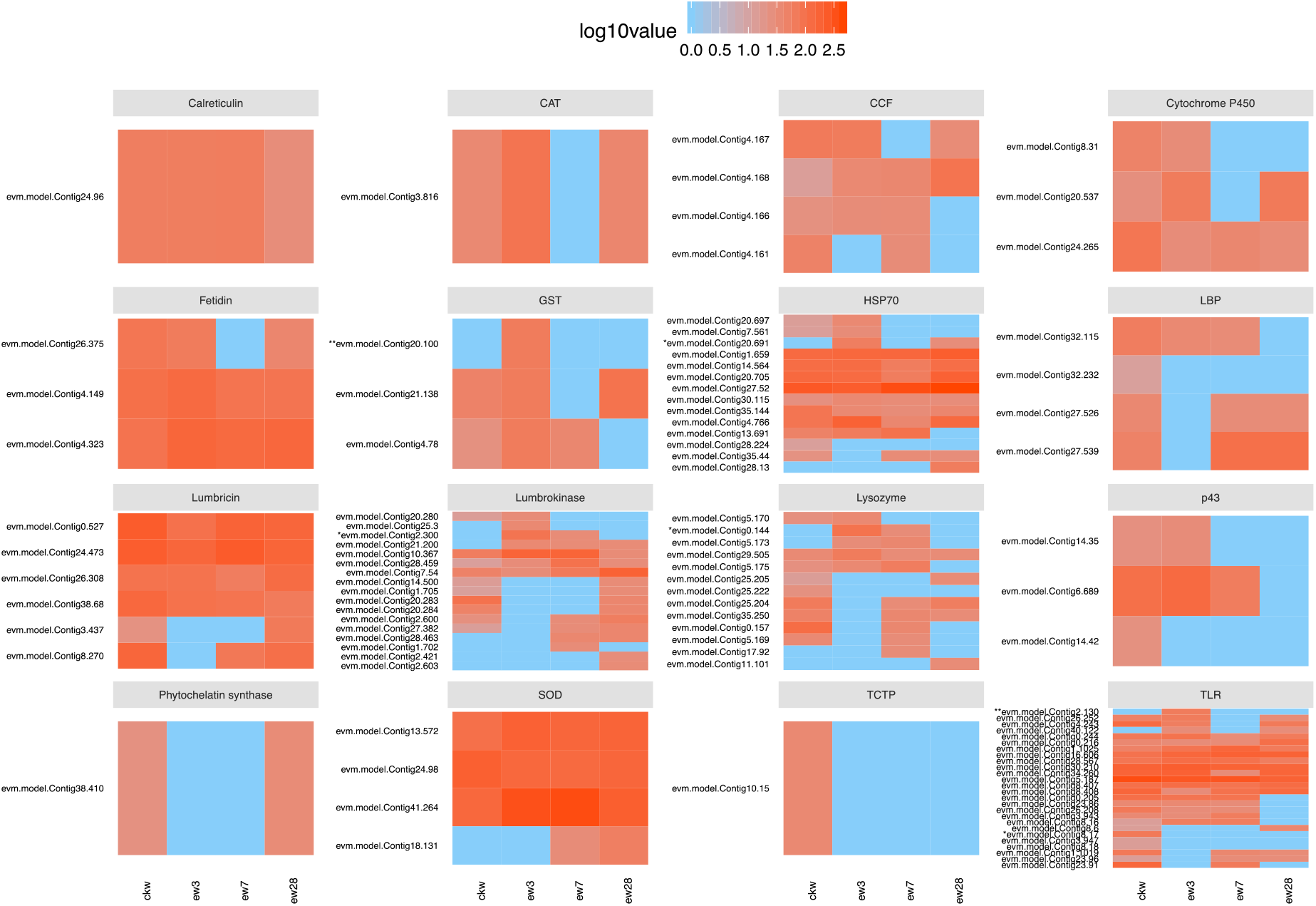
Expression heatmap of orthologous group members of well determined defensive genes in earthworm along days after feeding of pathogenic bacteria. In which, orthologous group members were clustered based on normalized expression count produced by protein mass spectrum.

## Discussion

Profiling of molecule mechanisms behind defensive of earthworm help to better understand the global distribution of this species and the origin of sophisticated immunity in vertebrates. In addition, it can also facilitate the usage of earthworm as an animal model in comparative immunology studies and ecotoxicology. Except for its value in research communities, earthworm is useful in monitoring environmental pollutions, and furthermore, is considered as an abundant source of biological active compounds with potential usage in industrial or medical fields.

## Supplementary Materials

**Supplementary Figure 1 | Genome assembly and annotation. a**, Heatmap of interaction intensity of intra-chromosome and inter-chromosome contacts detected by Hi-C for 42 chromosome-level scaffolds. **b**, The genome-wide distribution of annotated functional elements. The outermost circle represents 42 assembled chromosome-level scaffolds. Then, from outside to inside, the four circles represent the distribution of repeat sequences, non-coding genes, coding genes and SNPs, respectively. Connecting lines in the center represent duplicated genomic regions. The gray, blue and red colors of the connecting lines represent alignment scores generated by MCScanX that are lower than 500, between 500 and 1,000, and greater than 1,000, respectively.

**Supplementary Figure 2 | Odds ratios of significantly enriched functions related to the GO slim of immune system process along evolutionary steps.**

**Supplementary Figure 3 | Odds ratios of significantly enriched functions related to the GO slim of response to stress along evolutionary steps.**

**Supplementary Figure 4 | Odds ratios of significantly enriched functions related to the GO slim of homeostatic process along evolutionary steps.**

**Supplementary Table 1 Information of orthologous group members of well determined defensive genes in earthworm.**

## Acknowledgments

We thank NextOmics (Wuhan, China) for generating the PacBio and Hi-C data. We also thank Yan Sun, Qingxiao Li, and Qiong Yu for comments on the manuscript. We also thank Yanxia Hu, Xichun Zhang, Guozhen Xu, Yanqin Liu, Chong Wang, Xiaoping Diao, Yuhong Gao, Shuaizhang Li, Yanrui Luo, Xuelian Liu, Lan Yao, Feifan Guo, and so on, who are all previous members of Sun’s earthworm research laboratory.

## Author Contributions

X.W., Y.Zhao and Z.J.S designed and supervised research. X.W., Y.Zhang, Y.B.L. collected materials for sequencing and generated transcriptome data, proteomics, 16S rDNA and karyotype data. M.M.K. performed the genome assembly, genome annotation and heterozygosity estimation. Y.Zhao performed the synteny analysis, species tree, transcriptome, proteomics, 16S rDNA analysis. X.W. performed karyotype analysis. X.W., Y.Zhao, Y.Zhang, Y.F. Zhang, M.M.K, and Z.J.S. wrote the paper with assistance provided by co-authors.

---

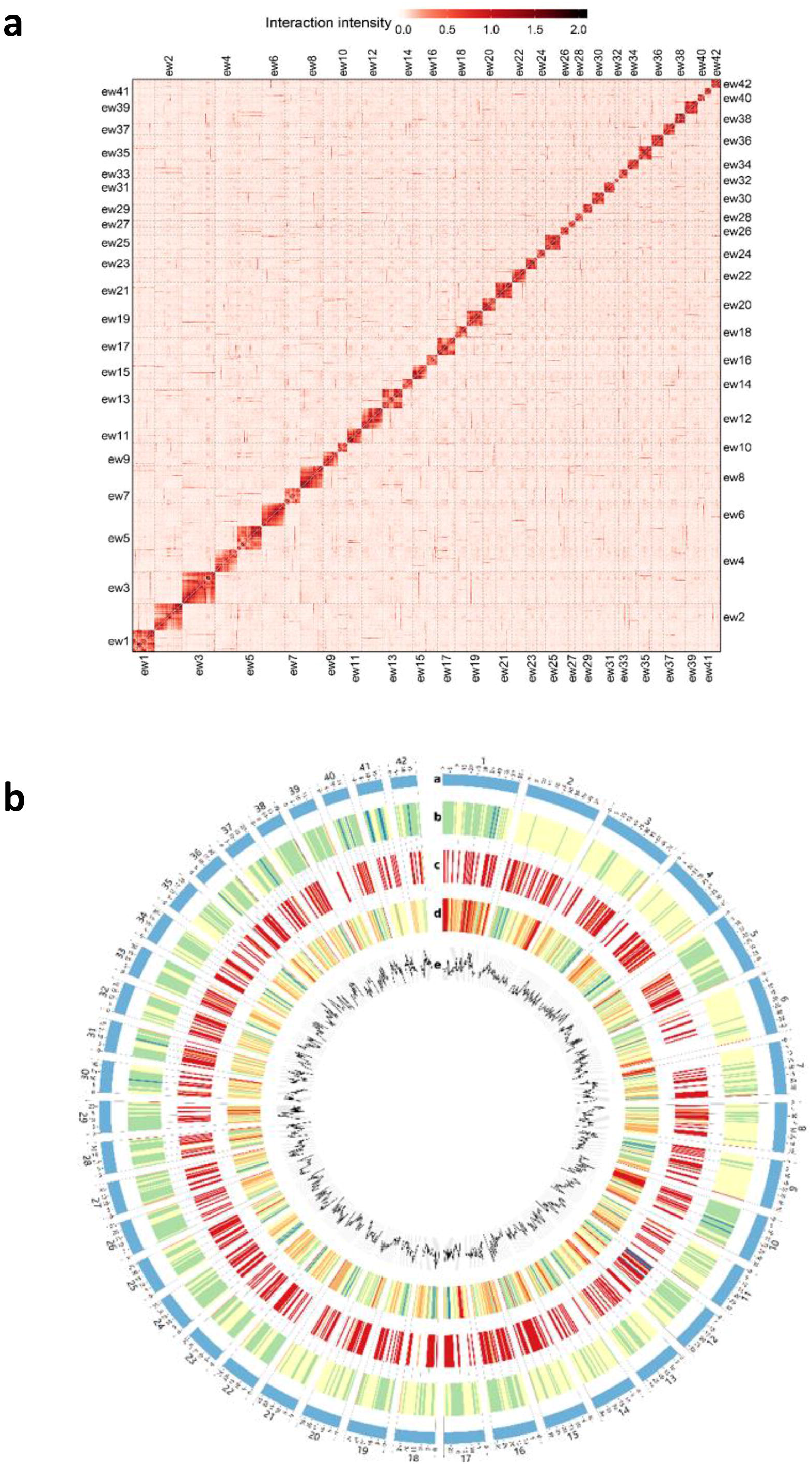

