## Supplementary figures and images for "Molecular mechanisms behind global distribution of earthworm revealed by the genome"

### Supplementary Figure 1

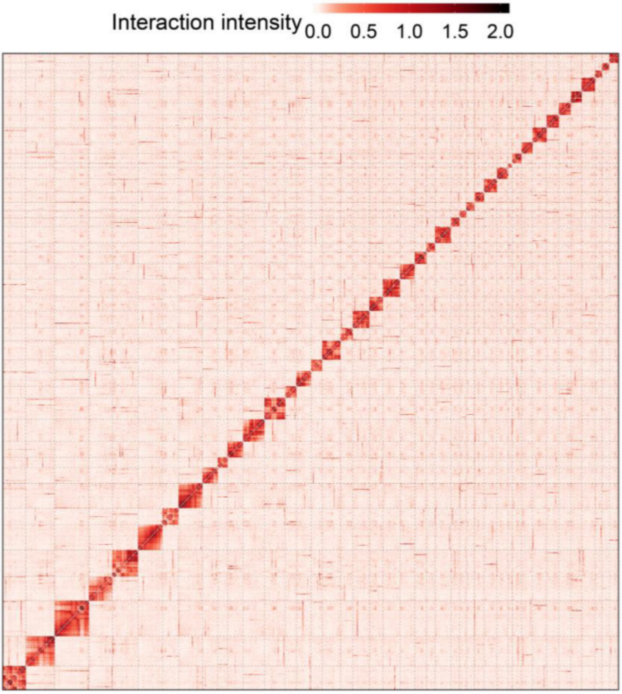

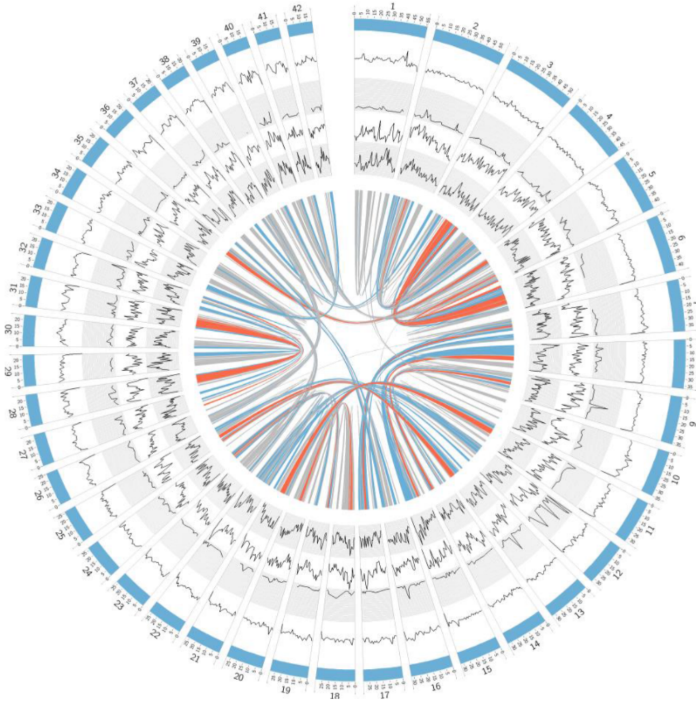

### Supplementary Figure 2

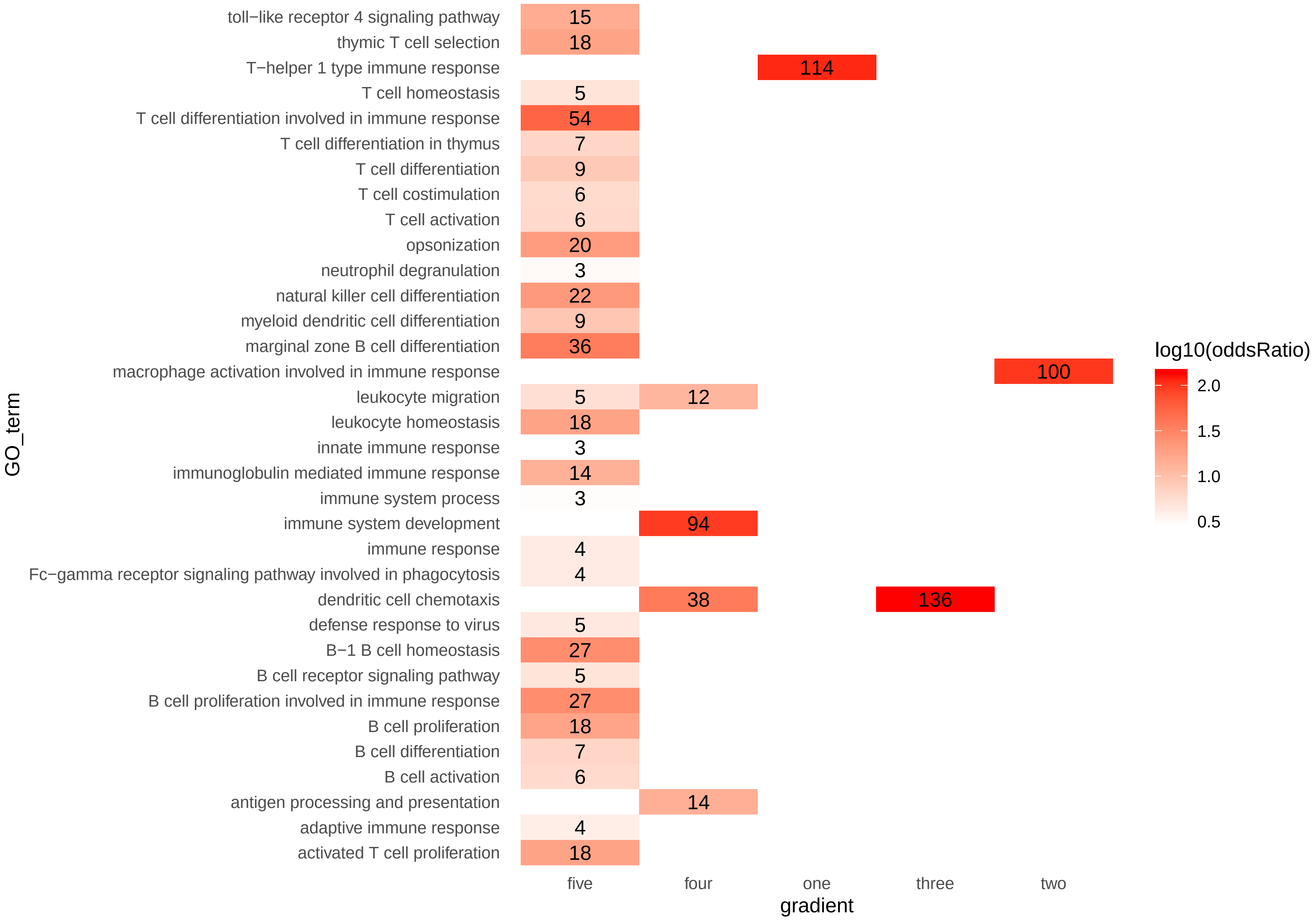

### Supplementary Figure 3

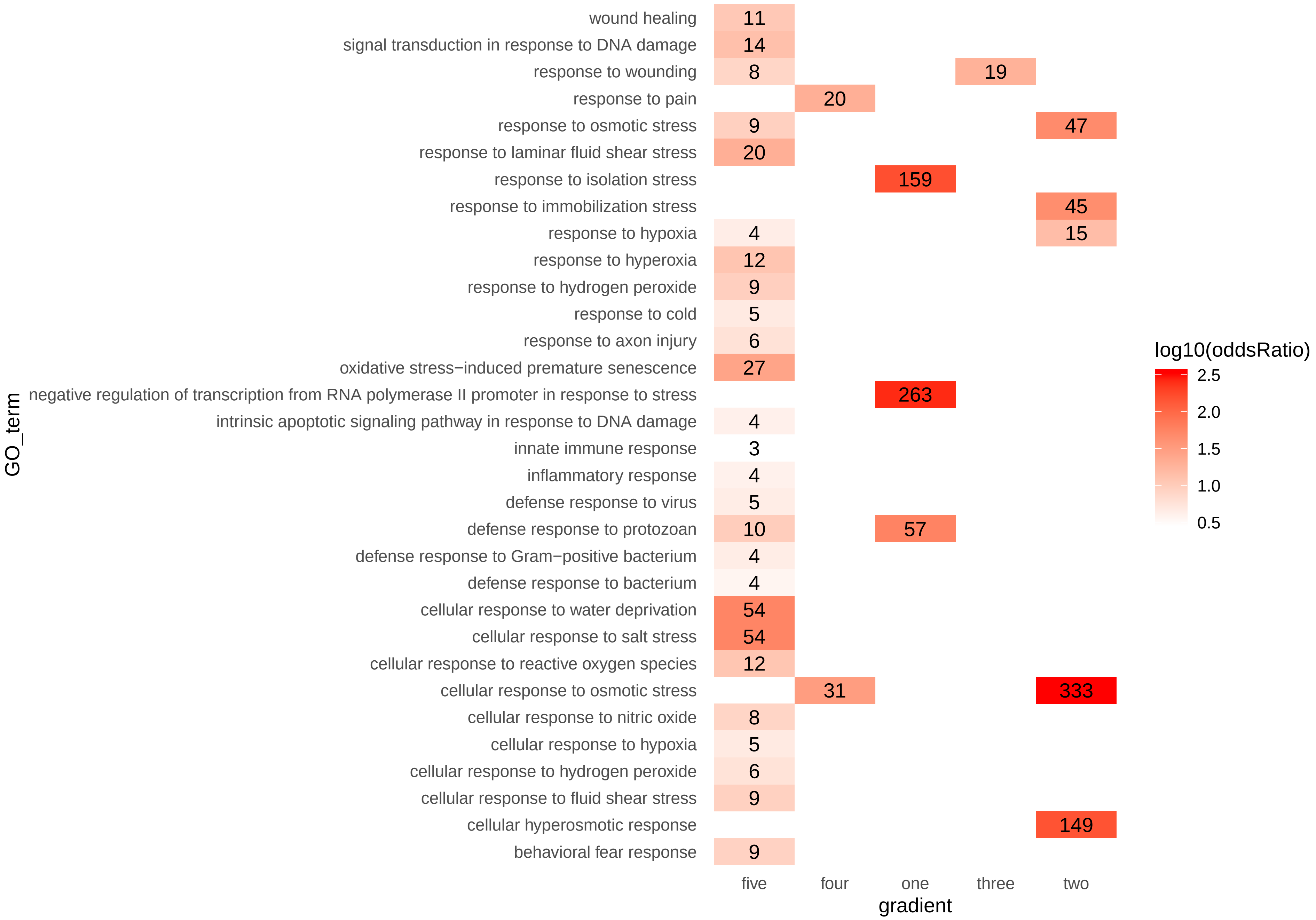

### Supplementary Figure 4

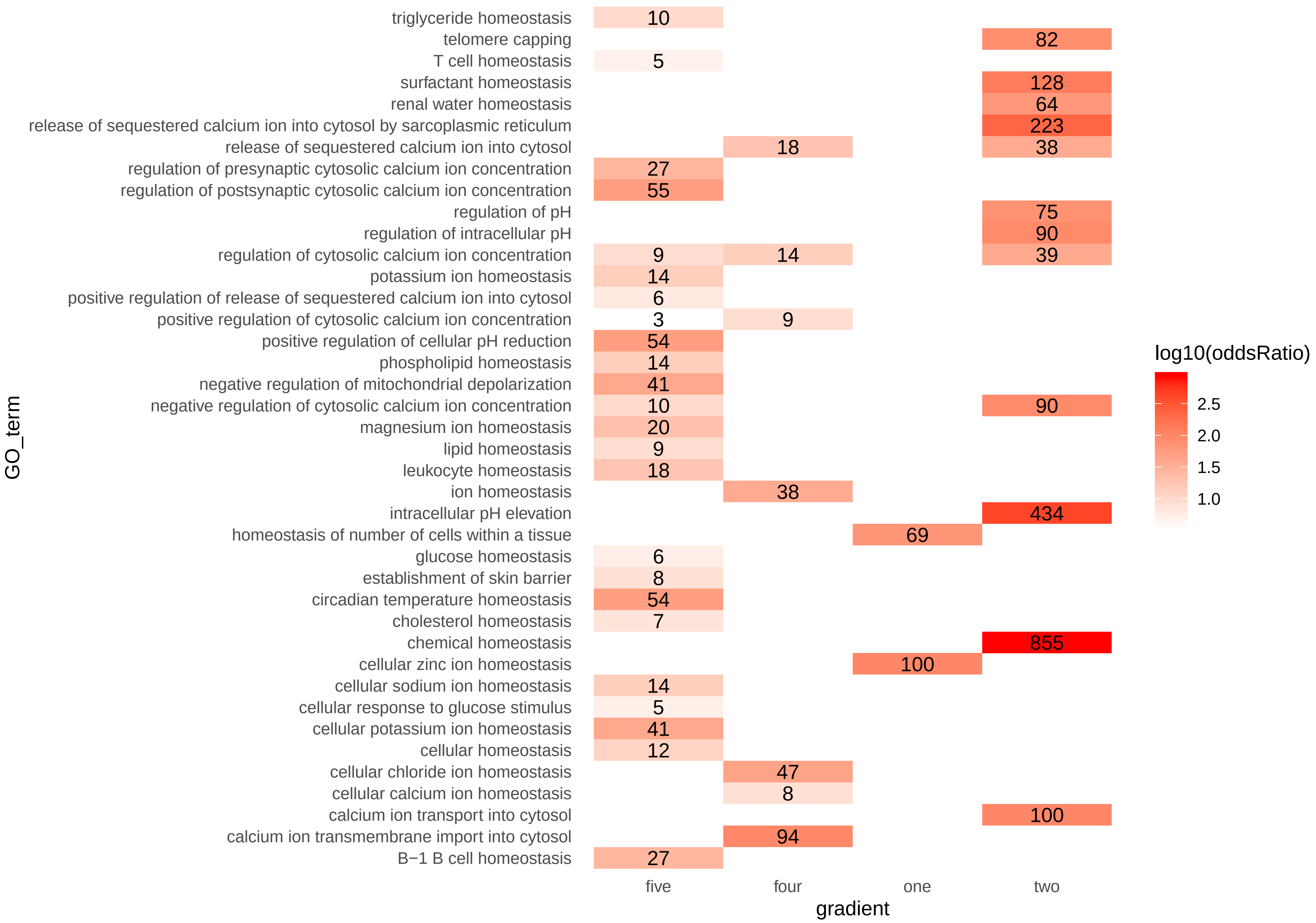
